## Supplementary material for "Bidirectional dysregulation of synaptic glutamate signaling after transient metabolic failure"

**Supplementary information**

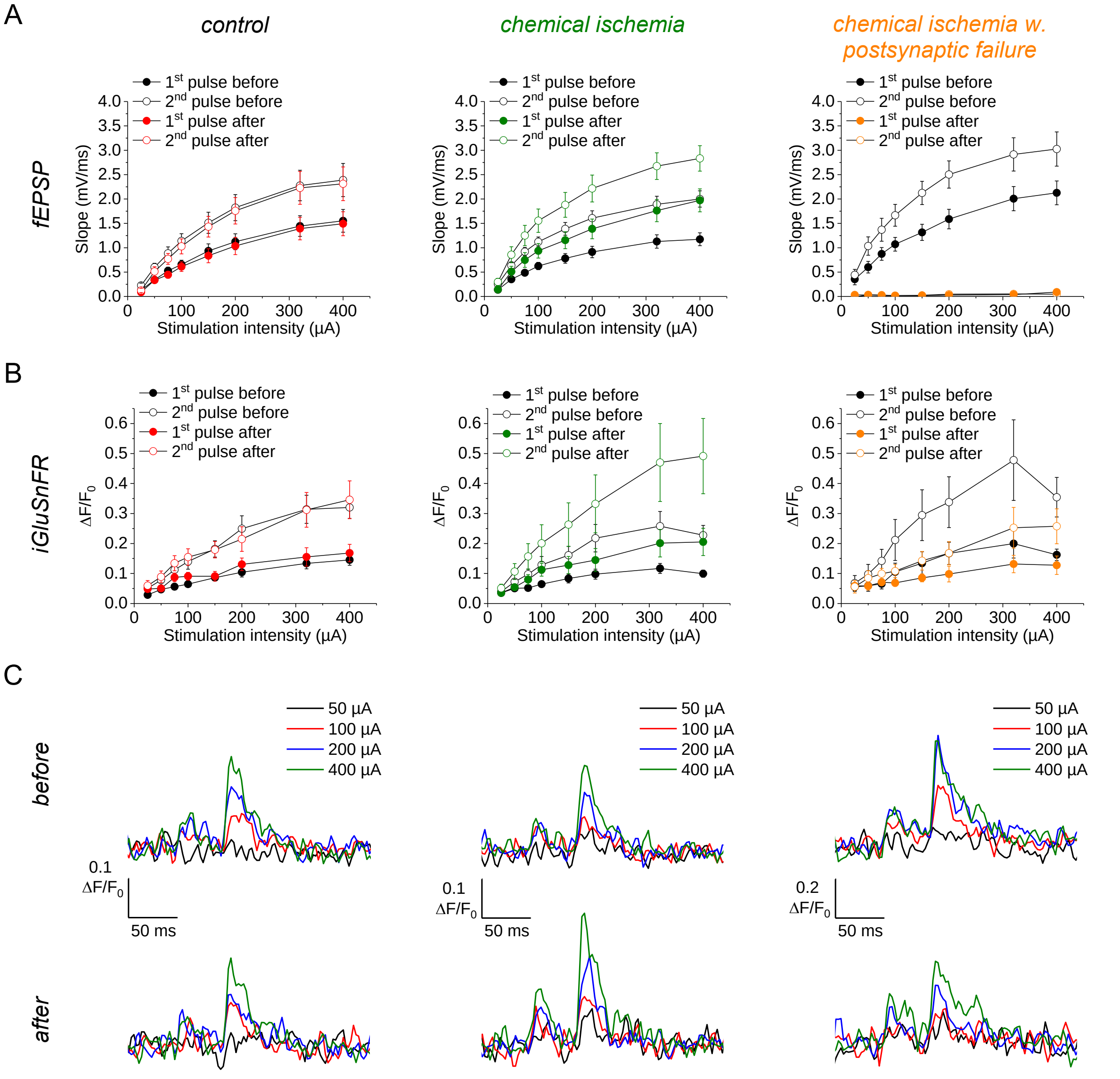


**Supplementary figure 1: Input-output relationship of fEPSP slope and iGluSnFR $\Delta F/F_0$ after chemical ischemia.** Further analyses of recordings presented in Fig. 2. (A) fEPSP slope in response to increasing stimulation intensities before and after control recordings (left, n = 8), recordings with chemical ischemia (middle, n = 9) and chemical ischemia with postsynaptic failure (right, n = 5). For each stimulation intensity two sweeps of a paired-pulse (50 ms, 1$^{st}$ and 2$^{nd}$ pulse) protocol were recorded. (B) Same as in (A) but for iGluSnFR $\Delta F/F_0$ amplitude in response to 1$^{st}$ and 2$^{nd}$ pulse. (C) Example traces of iGluSnFR $\Delta F/F_0$ recordings before (top row) and after (bottom row) control (left), chemical ischemia (middle) and chemical ischemia with postsynaptic failure (right) for 50, 100, 200 and 400 µA stimulation intensity (average of 2 scans/stimulation intensity). Data are expressed as mean ± s.e.m.

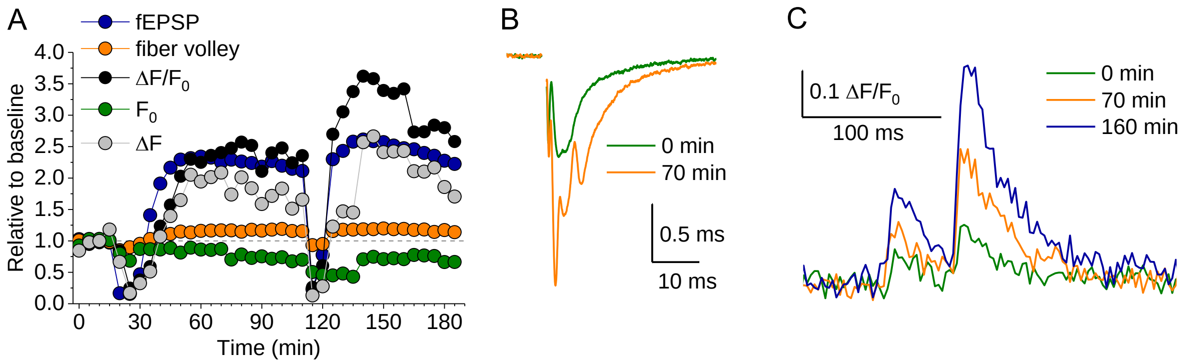


**Supplementary figure 2: Repeated chemical ischemia.** In a single experiment we have been able to induce acute chemical ischemia twice with stable electrophysiological and imaging for more than three hours. Note the persistent potentiation of the fEPSP slope and glutamate transients. (A) Time course of the indicated parameters (also see **Fig. 2**) including the resting iGluSnFR fluorescence before the stimulus ($F_0$) and its absolute change after the stimulus ($\Delta F$). The two dips at 20 min and 100 min indicate the two time points of chemical ischemia. (B) Example electrophysiological traces at the color-coded time points. (C) Example of iGluSnFR line scans at the color-coded time points.

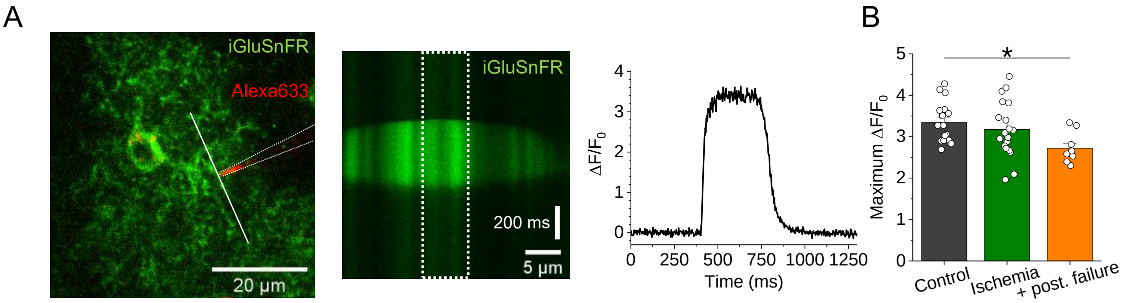


**Supplementary figure 3: Dynamic range of iGluSnFR is not affected by moderate chemical ischemia without postsynaptic failure.** (A) Left, example of an iGluSnFR-expressing astrocyte in the *stratum radiatum* of the CA1 region and iontophoresis micropipette (red, dotted line) filled with glutamate (150 mM) and Alexa Fluor 633 (40 µM). Middle, example of two-photon excitation line scan imaging (910 nm) of iGluSnFR fluorescence right in front of the iontophoresis micropipette tip in the periphery of an astrocyte (left, white line) during glutamate ejection by a 100 nA current (250 ms) pulse to saturate the iGluSnFR sensor. Right, change in iGluSnFR fluorescence ($\Delta F/F_0$) of line scan (middle, white dotted box). (B) Quantification of maximum $\Delta F/F_0$ amplitude during a saturating iontophoretic glutamate application pulse for control, chemical ischemia, and chemical ischemia with postsynaptic failure (+ post. failure). $p = 0.031$, one-way ANOVA; control vs. chemical ischemia, $p = 0.641$; control vs. chemical ischemia with postsynaptic failure. $p = 0.024$; chemical ischemia vs. chemical ischemia with postsynaptic failure, $p = 0.119$; $n = 18$, 19, and 9 from left to right; post-hoc Tukey test. Data are expressed as mean ± s.e.m.

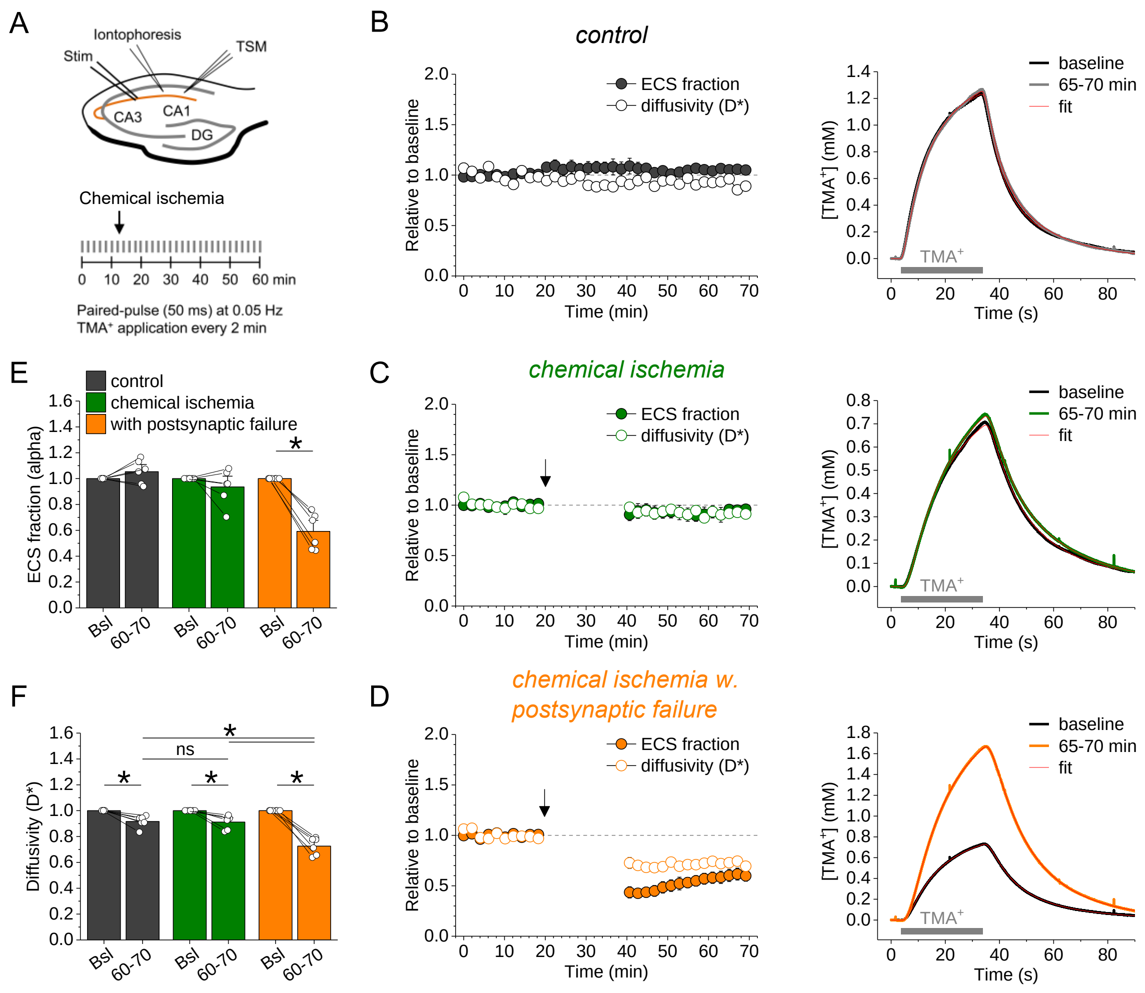


**Supplementary figure 4: The configuration of the extracellular space is only affected after severe chemical ischemia.** (A) Schematic of experimental design. Iontophoretically (Iontophoresis) applied $TMA^+$ transients (30 s every 2 min) were detected with a TMA-sensitive microelectrode (TSM) in the CA1 region during control recordings, and recordings with chemical ischemia without and with postsynaptic failure. Schaffer collaterals were stimulated as before (Stim; paired pulse (50 ms), 0.05 Hz). (B-D) Left, relative change of extracellular space (ECS) fraction and diffusivity (D*) compared to baseline (0-10 min) for control (B), chemical ischemia (C) and chemical ischemia with postsynaptic failure (D). Arrow indicates nominal start of chemical ischemia. Right, example traces of $TMA^+$ transients during baseline and last 5 min of recording (65-70 min) with corresponding fits (red). (E) Quantification of the ECS fraction in the last 10 min of recording (60-70 min) compared to baseline (Bsl). control: $p = 0.210$, $n = 6$; chemical ischemia: $p = 0.297$, $n = 6$; chemical ischemia with postsynaptic failure: $p = 0.0008$, $n = 6$; paired Student's t-test. (F) As in (E) but for diffusivity (D*). control: $p = 0.0063$; chemical ischemia: $p = 0.0105$; chemical ischemia with postsynaptic failure: $p = 0.0002$; paired Student's t-test. One-way ANOVA of effects after 60-70 min: $p < 0.0001$. Post-hoc Tukey tests: control 60-70 min vs. chemical ischemia 60-70 min, $p = 0.990$; control 60-70 min vs. chemical ischemia with postsynaptic failure 60-70 min,

$p < 0.0001$; chemical ischemia 60-70 min vs. chemical ischemia with postsynaptic failure 60-70 min, $p = 0.0001$. Data are expressed as mean ± s.e.m.

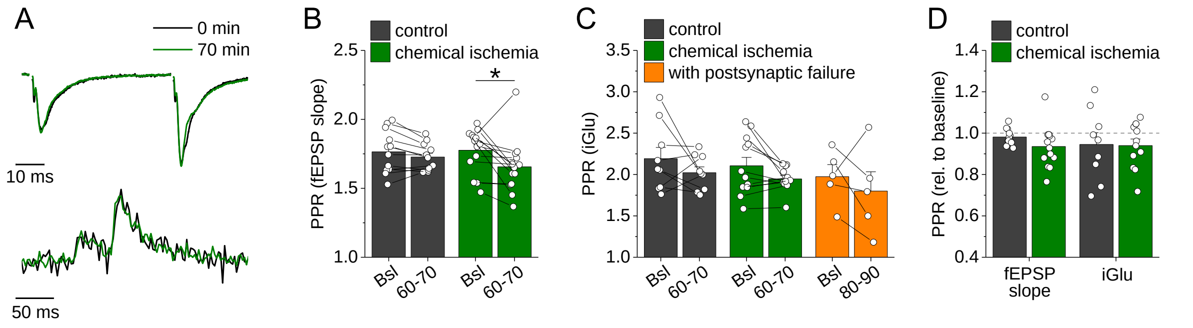


**Supplementary figure 5: Effect of transient chemical ischemia on short-term synaptic plasticity.** (A) Example traces of fEPSPs (top) and iGluSnFR $\Delta F/F_0$ fluorescence (bottom, average of 6 scans) before (black) and after (green) chemical ischemia without postsynaptic failure peak-scaled to baseline trace (same examples as in Fig. 2D but peak-scaled). (B) Quantification of paired-pulse ratio (PPR) of the fEPSP slope ($2^{nd}$ fEPSP slope / $1^{st}$ fEPSP slope) during the last 10 min of the recording (60-70 min) compared to baseline (Bsl) for control and chemical ischemia. Loss of the fEPSP in postsynaptic failure prevents analysis in these recordings. Control: p = 0.082, n = 12; chemical ischemia, p = 0.033, n = 14; paired Student's t-tests. (C) Same as in (B) but for PPR of iGluSnFR $\Delta F/F_0$ amplitudes. Control: p = 0.256, n = 9; chemical ischemia, p = 0.065, n = 12; chemical ischemia with postsynaptic failure, p = 0.454, n = 5; paired Student's t-tests. (D) Relative change of PPR compared to baseline for fEPSP slopes and iGluSnFR $\Delta F/F_0$ amplitudes for control and chemical ischemia. fEPSP slope: p = 0.127, n = 12 and 14; iGlu, p = 0.933, n = 9 and 12; Two-sample Student's t-test (with Welch correction for unequal variances for the fEPSP slopes). Data are expressed as mean ± s.e.m.

We conclude that there is no relevant change of short-term synaptic plasticity after chemical ischemia because the reduction of the electrophysiological PPR compared to baseline is not statistically different from those observed in control recordings (B, D) and there is no statistically significant change of the PPR of iGluSnFR transients (C, D).

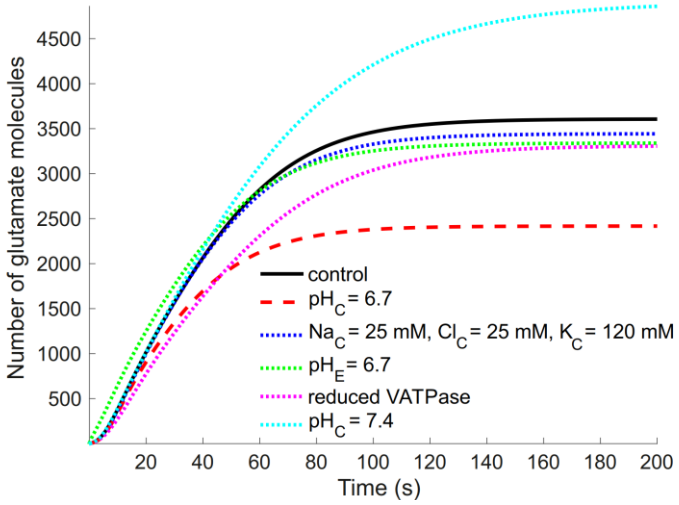


**Supplementary figure 6: Reduced glutamate concentration in synaptic vesicles in conditions typical of chemical ischemia.** Evolution of the number of glutamate molecules in synaptic vesicles under normal condition (control, black), reduced cytosolic pH ($pH_{C,}$ red), higher $Na^+$ and $Cl^-$ and lower $K^+$ concentration in the cytoplasm (blue), lower extracellular pH ($pH_E$, green), and reduced vesicular ATPase (VATPase) activity (pink). For illustration, the effect of a higher $pH_C$ is displayed (cyan), which is however not expected to occur in chemical ischemia.
